# A toolkit to handle T-less alignment for U insertion/deletion RNA editing

**DOI:** 10.1101/2022.07.06.498705

**Authors:** Fan Zhang, Ruslan Aphasizhev, Inna Afasizheva, Liye Zhang

**Affiliations:** Shanghaitech; Boston University Medical Campus; ShanghaiTech University

## Abstract

**Summary:** In the mitochondrial genome of kinetoplastids, the targeted insertion and deletion of uridine in cryptic mRNA is essential to generate functional protein-coding mRNAs, but such process leads to a highly hetergenous and complex RNA population. Few tools were developped to perform alignment based on a T depleted strategy for such mRNA population, however to our knowledge no downstream analysis tool is available so far to allow easy data processing and interpretation. Here, we fill this gap with the *T-less-Alignment-Toolkit*, which provides useful functions such as duplication removal with UMI, alignment strandness analysis, statistical summary and visualization.

**Availability and Implementation:** Scripts and toolkits can be downloaded at https://gitee.com/Zhanglab/tless-alignment-toolkit

## 1 Introduction

Mitochondrial DNA (mtDNA) of kinetoplastid protist contains thousands of interlinked circular molecules arranged in a disk-like network. There are two types of circular molecules called minicircles and maxicircles. Minicircles encode various guide RNAs (gRNA) which mediate the U insertion/deletion editing (U-indel editing) of the pre-matured mRNAs encoded by maxicircles to make them functioned properly. (Aphasizhev R. and Aphasizheva I. 2011)

Study of this complex U-indel editing requires unique read mapping algorithm. Currently, tools like PARERS, TREAT, and *T-Aligner* have been constructed to analysis the U-indel system, but only *T-Aligner* can do the U-indel mapping with short reads sequencing. *T-Aligner* was written in C++ and managed to tackle U-indel editing alignment by masking all Ts and then perform 3-letter A/C/G alignment. (Gerasimov, E. S. et al. 2018)

However, we found that current version of *T-Aligner* misses some useful features such as handling PCR bias by UMI, matching raw sequenced reads to alignment output and extracting mapping strandness information. Here, we present a python toolkit, *T-less-Alignment-Toolkit*, which facilitates the downstream analysis of U-insertion/deletion mRNA profiles by *T-Aligner*.

## 2 Method

Program *alignlib* within *T-Aligner3*.*3* package was used to perform the T-less read alignment. Unlike the commonly used SAM file, the alignment output tar file only contains mapping information (See supplementary Fig S1 for a short introduction), but does not contain the original read information. Therefore, *T-less-Alignment-Toolkit* first takes the taf file given by *alignlib* as well as original FASTQ file as inputs to match the alignment output to original nucleotide sequences.

**Fig1.**
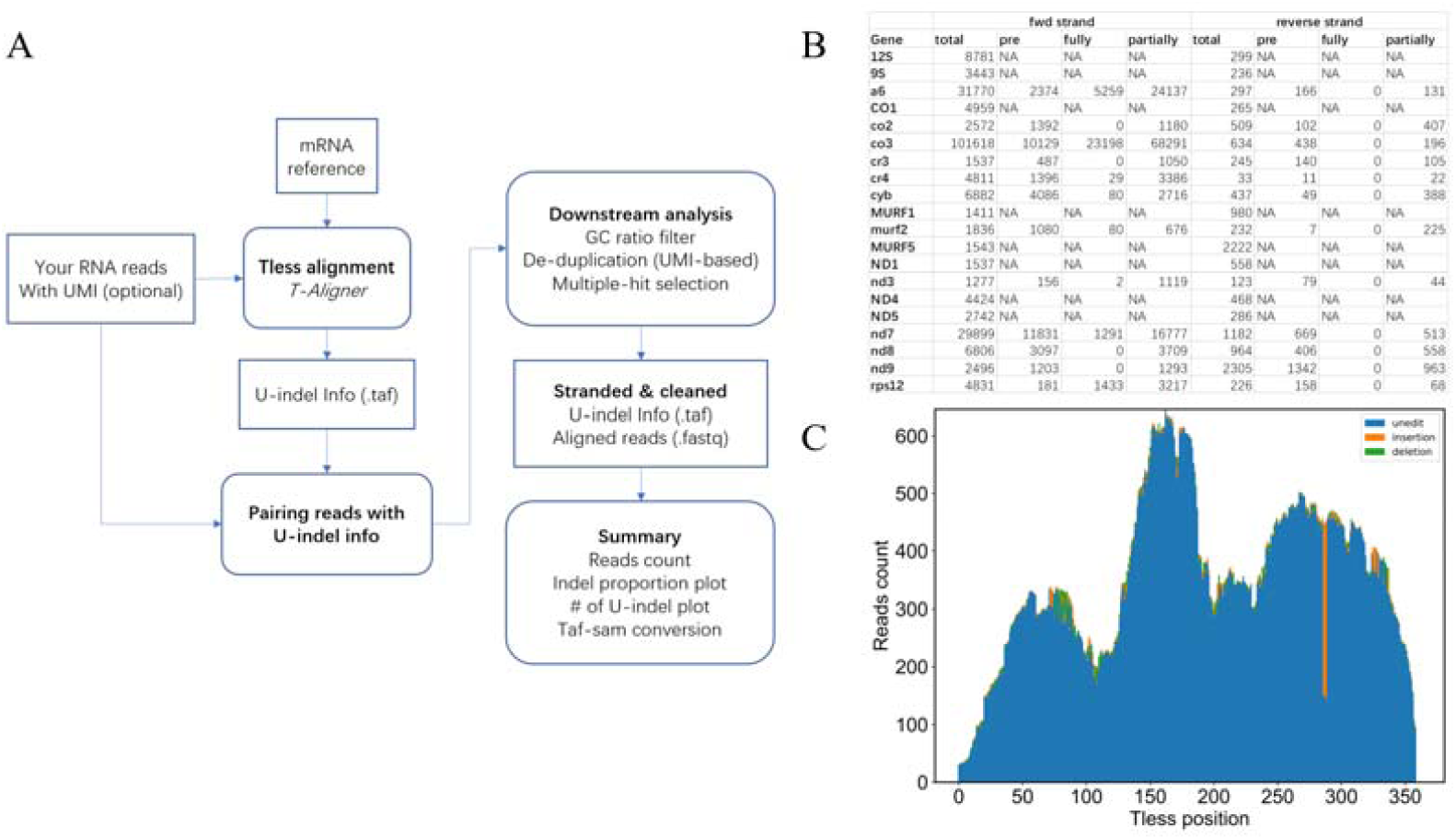
Pipeline of *T-less-Alignment-Toolkit*. (A) *T-less-Alignment-Toolkit* workflow. *T-less-Alignment-Toolkit* requires mainly two input data: RNA-seq reads and pre-edit mRNA reference. The toolkit will firstly use T-Aligner to do the alignment, followed with customed scripts to do downstream polish and analysis. (B) Statistical summary of eCLIP data for KREH2 protein (Van Nostrand, E. L. et al. 2016). For 12 cryptogenes (capital letter), genes that need to be edited, reads editing state will be given. While for 8 never-edit genes (lower case letter), genes that do not need to be edited, only total reads count for forward/reverse strand will be given. (C) Forward strand’s editing plot for co2 gene.

### 2.1 Pairing raw reads with U-indel information

*T-less-Alignment-Toolkit* will use the index (2^nd^ column) given by taf file to match the alignment information to the raw read. Since the index given by *alignlib* have a maximum number of 5,999,999, all input read library will be split into sub library with 2,500,000 lines each.

### 2.2 GC ratio filter and Read identification

Because T-Aligner use 3-letter sequence to do alignment, it has a poor performance when it encounters AT-rich region, showing abnormal spike in (Fig S2A). Besides, after using a moving window of 20nt for both edited and pre-edited RNA, we found that the cases with <20% CG is very small (Fig S2B). Thus, we set the threshold that the matched sequence (6^th^ column) with GC ratio < 20% will be discarded. After GC ratio filter, the normal pattern remains almost the same (Fig S2C).

Then, the ‘editing distance score’ between the raw read sequence and the forward/reverse strand of the reference sequence calculated by *python-Levenshtein* package will be used to identify the strandness of the read.

### 2.3 Remove duplication (optional)

For reads with the same UMI annotated by *UMI-tools* (Fig S3) and the same mapping condition (including start & end position and the editing matrix) given by *alignlib* are grouped together. For each group: (1) If all the sequence are the same, only one read will be retained; (2) If hidden SNP exists, for each SNP, the read that all SNP sites have the most frequent nucleotide will be retained; (3) Otherwise, the group will be discarded.

### 2.4 Multiple-hit selection

*T-Aligner* also encounters one read align to different genes. For these reads: (1) The longest alignment will be retained; (2) If reads have the same aligned length, reads with the least SNP against the reference will be retained; (3) Otherwise, all reads will be retained.

### 2.5 Further analysis

Apart from optimized T-less alignment, *T-less-Alignment-Toolkit* also provides scripts for further analysis and visualization of the U-indel editing.

#### (1) Summary part

First, the read counts for forward/reverse will be given. Besides, the editing state of the read will be counted as ‘pre edit’, ‘partially edited’ and ‘fully edited’.

To determine the editing state, reference number of T that should be inserted/deleted for each T-less site is calculated by matching pre edit reference against fully edited reference. For reads with no T editing, they are termed as ‘pre edit’. For reads with all the same number of T editing as the fully edited reference does, they are termed as ‘fully edited’. While the other reads are termed as ‘partially edited’.

#### (2) Plotting

The overall U-indel proportion plot per site will be plotted (Fig S4). Comparing with Fig S2C, the decline of reads count indicating the effect of duplication removal. After strandness identification, the different read distribution pattern between forward and reverse strand provided a new insight into U-indel editing analysis.

## 3 Conclusion

With *T-less-Alignment-Toolkit*, we managed to perform more precise and detailed U-indel editing analysis for UMI labeled (or not) datasets. This package may give a new insight into U-indel editing and can be easily used to analysis U-indel editing related network by conditional knockout.

## Supporting information

All

## Notes

https://gitee.com/Zhanglab/tless-alignment-toolkit/tree/master

### Competing Interest Statement

The authors have declared no competing interest.

### Summary of Updates

Update Main figure (Fig 1) with some example results. Update Reference format for Bioinformatics journal. Update supplementary figure (Fig S2A), specifying the reference sequence state.

