## Supplementary material for "A toolkit to handle T-less alignment for U insertion/deletion RNA editing": All

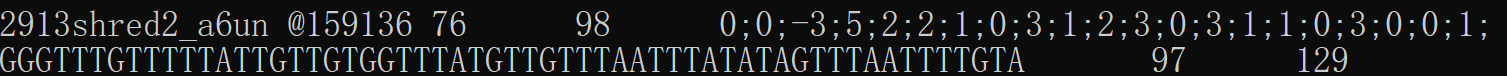

**Fig S1. Sketch of *alignlib* result taf file.** taf file output by *alignlib* is a regular text table, each line describes read mapping. There are 8 columns each line, 1^st^ column shows the name of reference sequence; 2^nd^ column gives the index of the read; 3-4^th^ columns show the start and the end position of the T-less reference; 5^th^ column gives the number of U-indels between two T-less letters; 6^th^ column gives the matched sequence in reference and 7-8^th^ column give the start and end match position of the reference.

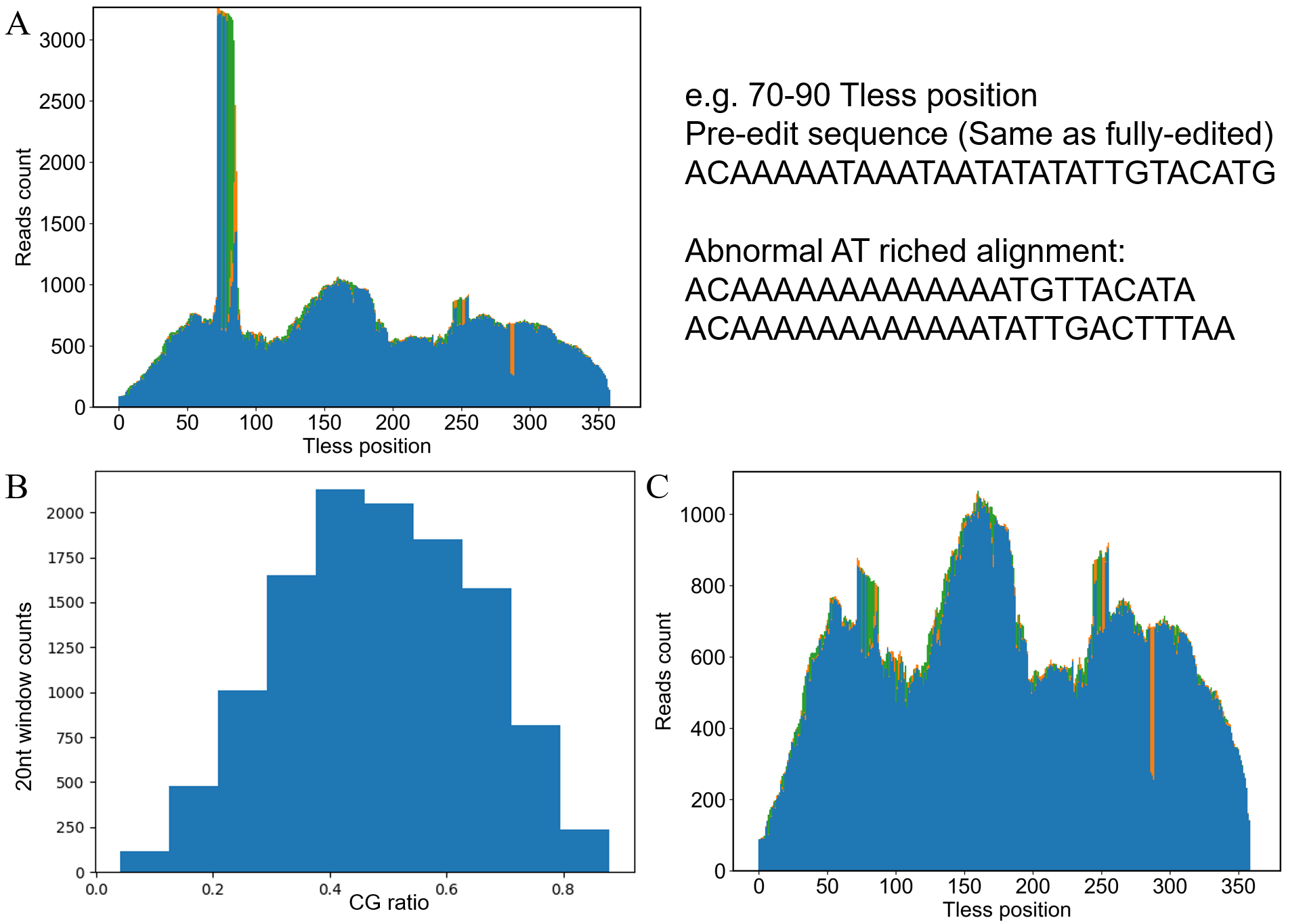

**Fig S2. High AT contained spike in and CG ratio distribution for both edited and pre-edit RNA.** (A) Editing plot (Green bars refer to deletion, orange bars refer to insertion and blue bars refer to non-edit) for co2 gene with eCLIP data of KREH2 protein before CG ratio filter. There is a strong spike in with strong abnormal editing at around 75 Tless position. The pre-edit sequence and some abnormal AT-rich raw reads are listed in the right. Notice that the 70-90 Tless position requires no U indel editing, so it is the same for pre-edit and fully-edit sequence. (B) A 20nt moving window constructed from both pre-edit and fully edited 12 cryptogenes. (C) Editing plot for co2 gene with eCLIP data of KREH2 protein after CG ratio filter. The strong spike in was mostly eliminated and the overall read distribution pattern remains and is much clearer.

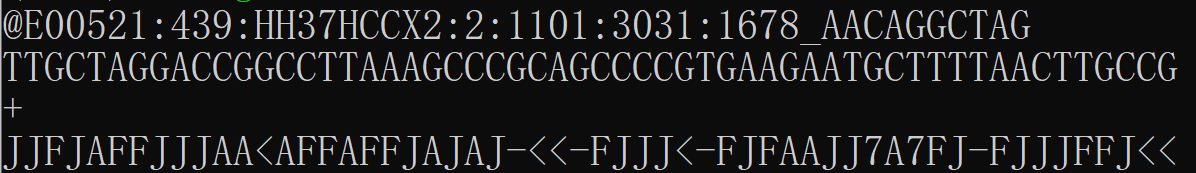

**Fig S3. Example of sequencing read labeled with UMI by UMI-tools (Red frame labeled).** The UMI was labeled at the end of the read name with ‘_’ connecting read name and the UMI sequence.

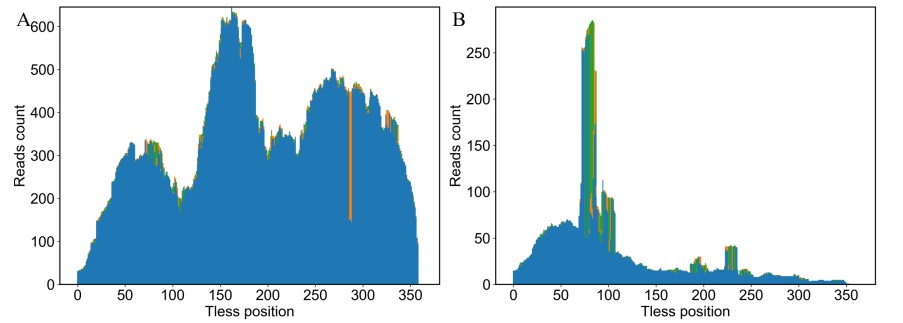

**Fig S4. Example of overall U-indel proportion plot.** (A) overall U-indel proportion plot for forward strand. (B) Overall U-indel proportion plot for reverse strand
